# Multi-omics fine-mapping uncovers a pleiotropic intestinal enhancer variant linking PMAIP1 regulation to pig growth and fatness

**DOI:** 10.64898/2026.09.26.754598

**Authors:** Hui Yang, Mingyang Hu, Yuanfeng Xu, Jinfu Zhang, Yonghui Zhang, Yuwei Gou, Xiaoyong Du, Heng Wang, Xinyun Li, Yunxia Zhao, Tao Xiang

## Abstract

Genetic loci influencing multiple complex traits often remain difficult to interpret because the underlying functional variants, target genes and tissue contexts are unresolved. Here, we investigated the regulatory basis of a shared signal for average daily gain (ADG) and backfat thickness (BF) in Yorkshire pigs. By integrating GWAS in 7,972 pigs, targeted capture sequencing, LD-based fine-mapping, porcine epigenomic annotations, three-dimensional chromatin interaction maps and functional assays, we narrowed the shared SSC1 signal to a 1.59-Mb interval and functionally prioritized rs334769214 (T>G) as the leading candidate variant. The T>G substitution produced marked allele-specific differences in enhancer activity and nuclear-protein binding, while transcription-factor co-transfection assays identified FOXL1-responsive, allele-dependent regulatory activity. Chromatin interaction and CRISPRa analyses supported a regulatory connection between the rs334769214-containing enhancer and PMAIP1, and PMAIP1 overexpression promoted apoptosis in porcine intestinal epithelial cells. In the capture-sequenced subset, the G allele was associated with higher ADG and BF, consistent with a relative loss-of-activation model. Together, these findings delineate a candidate FOXL1-enhancer-PMAIP1 regulatory pathway and demonstrate how multi-omics fine-mapping can advance a shared quantitative-trait locus toward a testable molecular mechanism underlying variation in growth and fatness.

## Introduction

Growth rate and fat deposition are fundamental quantitative traits that reflect how nutrients are partitioned between lean tissue accretion and energy storage in pigs. Average daily gain (ADG) captures postnatal growth performance and production efficiency, whereas backfat thickness (BF) reflects subcutaneous fat deposition and carcass composition [1–2]. Although these traits are routinely considered in breeding programs, they also provide a useful biological model for studying the genetic regulation of growth-fatness balance. Both ADG and BF are heritable, and previous studies have reported variable but often positive genetic correlations between growth rate and fat deposition across pig populations [2–5]. This relationship suggests that some genomic regions may influence the two traits in a coordinated manner, although the underlying molecular mechanisms remain poorly understood.

Genome-wide association studies (GWAS) have become a powerful approach for mapping quantitative trait loci (QTLs) associated with complex traits [6]. In pigs, thousands of QTLs for ADG, BF and related production traits have been cataloged in PigQTLdb [7]. However, most GWAS signals remain statistical associations. For many loci, especially those located in non-coding regions, the functional variant, effector gene, relevant tissue context and regulatory mechanism remain unresolved [8]. This limitation becomes more pronounced when the same genomic region is associated with multiple traits, because association data alone cannot distinguish whether the observed signals arise from a shared functional variant, linked variants in linkage disequilibrium or independent regulatory mechanisms.

Functional genomics and three-dimensional chromatin datasets provide a route from associated loci to testable regulatory mechanisms. Chromatin accessibility, histone modification profiles and transcription factor-binding annotations can help prioritize non-coding variants with regulatory potential, whereas chromatin conformation data can connect distal enhancers with their putative target genes [9–11]. In livestock, recent functional annotation and multi-omics resources, including porcine epigenomic maps and PigGTEx-informed analyses linking variants to genes and traits, have substantially improved the interpretation of complex-trait loci [12–14]. Nevertheless, few studies have resolved the functional variant, target gene, tissue context and transcription factor logic underlying genomic regions jointly associated with ADG and BF.

Here, we combined large-scale GWAS, targeted capture sequencing, fine-mapping, porcine epigenomic annotation, 3D chromatin interaction analysis and functional assays in Yorkshire pigs to characterize the regulatory basis of a shared SSC1 association signal for ADG and BF. By integrating population-level association, regulatory annotation, enhancer assays, EMSA, CRISPR activation and cellular phenotyping, we prioritized an intestinal enhancer variant linked to PMAIP1 regulation. Our study provides an integrative framework for prioritizing candidate non-coding regulatory variants and linking them to potential target genes through allele-specific regulatory activity and enhancer-promoter interactions underlying variation in growth and fat deposition.

## Materials and methods

### Ethics statement

All animal procedures and tissue collections were approved by the Ethics Committee of Huazhong Agricultural University (approval NO. HZAUSW-2018-008) and were conducted in accordance with institutional animal care guidelines.

### Animals, phenotypes and genotyping

The study population comprised Canadian Yorkshire pigs raised under standardized farm management. BF was measured between the 10th and 11th ribs at a body weight of 100 ± 15 kg using an Aloka 500V SSD B-mode ultrasound instrument. The birth-to-100-kg average daily gain (ADG) and backfat thickness (BF) records were collected from March 2014 to January 2018, yielding 58,759 ADG records and 58,736 BF records. The pedigree database contained approximately 640,000 records and enabled pedigree tracing for more than 5 generations. For genomic analyses, animals with both phenotypic records and ear tissue samples were retained, including 7,972 pigs for ADG and 7,959 pigs for BF. Genotyping was performed using the Illumina Customer-designed PorcineSNP60 BeadChip array. The raw dataset contained 53,261 autosomal SNPs, which were mapped to the Sscrofa11.1 reference genome assembly. Genotype quality control was performed using the following thresholds: individual call rate >= 90%, SNP call rate >= 90%, minor allele frequency >= 0.01 and Hardy-Weinberg equilibrium P >= 1 x 10^-6^. After filtering, 41,843 high-quality SNPs were retained for ADG and 41,845 for BF. The detailed quality control can be found in our previous study [15]. These filtered datasets were used for all subsequent association analyses.

### Phenotype pre-correction

Phenotypic records for ADG and BF were pre-corrected using the DMU software package [16].

Variance components and genomic estimated breeding values were estimated using single-step genomic best linear unbiased prediction, which integrates pedigree and genomic information [17–18]. The following linear mixed model was fitted:

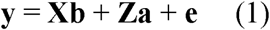

where **y** is the vector of original phenotypic values; **b** is the vector of fixed effects, including herd-year-season, sex and age; **X** is the incidence matrix for fixed effects; a is the vector of random additive genetic effects; **Z** is the incidence matrix for additive genetic effects; and **e** is the vector of residuals. Residual effects were assumed to follow e ∼ N(0, **I**σ^2^*_e_*), where I is the corresponding identity matrix and σ^2^*_e_* is the residual variance; additive genetic effects a were assumed to follow a ∼ N(0, **H**σ^2^*_a_*), where **H** is the relationship matrix combining pedigree and genomic information [17], and σ^2^_u_ is the genetic variance. Corrected phenotypes Y_c_ for both traits were calculated as the sum of the estimated additive genetic effects and residuals:

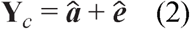

where **Y**_c_ is the corrected phenotype, a is the estimated additive genetic effect and e is the residual term.

### Genome-wide association analysis

GWAS for ADG and BF was performed using rMVP [19]. The analysis included 7,972 genotyped pigs for ADG and 7,959 genotyped pigs for BF. A mixed linear model was used:

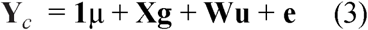

where Y_c_ is the vector of corrected phenotypes; μ is the overall mean of corrected phenotypes; **1** is a vector of ones; **X** is the genotype matrix coded as 0, 1 or 2 for the three SNP genotypes; **g** is the fixed additive effect of the tested SNP; **u** is the vector of random polygenic effects, assumed to follow **u** ∼ N(0, **G**σ^2^_u_), where **G** is the marker-based additive genomic relationship matrix constructed as described previously [24], and σ^2^_u_ is the polygenic variance; W links individuals to the polygenic effects; and e is the residual vector, assumed to follow e ∼ N(0, **I**σ^2^_e_), where σ^2^_e_ is the residual variance. Genome-wide significance was determined using Bonferroni correction, with the threshold calculated as −log10(0.05/N), where N is the number of SNPs tested. This corresponded to a genome-wide threshold of 5.93 for the SNP array analysis.

### Haplotype and QTL mapping analysis

LD blocks surrounding SNPs significantly associated with ADG or BF were defined using Haploview with the confidence interval algorithm [20]. QTLs located within these LD blocks were retrieved from PigQTLdb (https://www.animalgenome.org/cgi-bin/QTLdb/SS/index) [7]. Haplotype construction on Sus scrofa chromosome 1 (SSC1) was performed using the LD and linkage analysis framework [21–22]. This approach combines population-level LD with within-family linkage information to improve QTL localization [21–25]. Haplotypes were constructed using PHASEBOOK [23] under a hidden Markov model, assuming 20 ancestral haplotypes. QTL mapping was performed using a likelihood ratio test that compared a model including haplotype effects with a reduced model excluding haplotype effects [26]. The model including haplotype effects was:

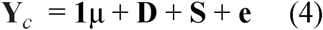

where **D** and **S** represent random haplotype effects inherited from dam and sire, respectively, with **D** ∼ N(0, **I**σ^2^_hd_) and **S** ∼ N(0, **I**σ^2^_hs_), where σ^2^_hd_ and σ^2^_hs_ are variances for haplotype effects in dam and sire, respectively. A 95% confidence interval for QTL position was defined as the region in which the likelihood ratio test statistic remained within 2 units of its maximum value [26].

### Targeted capture sequencing and association analysis

To refine the candidate interval, targeted deep sequencing (>20x coverage) was performed for a 6.37 Mb interval on SSC1 (159.49-165.86 Mb) in 872 randomly selected genotyped individuals. Custom targeted-capture libraries were constructed from isolated genomic DNA using a probe-based target enrichment protocol [27]. Qualified libraries were sequenced on the Illumina HiSeq3000 platform to generate paired-end reads. Raw reads were quality filtered with Trimmomatic v0.33.0 and aligned to the Sscrofa11.1 reference genome using BWA v0.7.15 [28]. Variant calling was performed using GATK4 [29]. SNPs were excluded if they met any of the following criteria: QUAL < 30.0, QD < 2.0, ReadPosRankSum < −8.0 or FS > 60.0. Only variants marked as “PASS” were retained for subsequent analyses. Association analysis of sequence-derived SNPs was performed for variants within the refined candidate intervals: 159.84-162.41 Mb for ADG and 159.49-165.86 Mb for BF. The following mixed linear model was fitted in rMVP [19]:

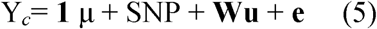

where SNP represents the fixed effect of the tested sequence variant with three genotype classes, and the remaining terms are as defined above. To reduce over-correction caused by including variants from the tested region in the relationship matrix, the genomic relationship matrix was constructed using a leave-one-chromosome-out strategy that excluded SSC1 [30]. Multiple testing was controlled using Bonferroni correction with the threshold calculated as −log10(0.05/N), where N is the number of SNPs tested.

### Integration of porcine epigenomic datasets

Porcine epigenomic datasets based on Sscrofa11.1, including topologically associating domains, Hi-C contact matrices, H3K27ac peaks and enhancer-gene pairs, were obtained from in-house porcine epigenomic datasets generated by our group. Hi-C reads were processed using HiC-Pro v2.9.0 [31]. Topologically associating domains (TADs) and subdomains were identified using insulation score and top-domain approaches [32]. For ChIP-seq analysis, reads were aligned with BWA v0.7.15, low-quality reads were filtered using SAMtools v1.9 and Picard v1.126, and peaks were called using MACS2 v2.1.0. Detailed procedures for porcine Hi-C and ChIP-seq processing were described previously [11]. GWAS signals were integrated with Hi-C contact maps and TAD structures using Juicebox v2.13.07 [33]. H3K27ac peaks across tissues were merged using BEDTools v2.26.0 [34], and regulatory signals around candidate genes were visualized with igv.js v3.1.3 [35].

### Prioritization of functional SNPs and target genes

Significant SNPs from the capture sequencing association analysis were submitted to IFmut (Integrated Functional Mutation database: http://119.96.199.105:8212/#/home) and prioritized using its functional scoring system [36]. Candidate variants classified as high-confidence regulatory variants (classes 1a-3b) were retained for motifbreakR analysis. Variants predicted to have a strong effect on transcription factor binding were selected for experimental validation. To identify potential target genes of prioritized functional variants, Pol II ChIA-PET chromatin interaction data generated from 12 Yorkshire pig tissues, including adipose, muscle and small intestine, were analyzed. Chromatin loops were identified using CPU v0.0.1 (https://github.com/cheehongsg/CPU) with reads aligned to the Sscrofa11.1 reference genome. Candidate SNPs were assigned to potential target genes by intersecting SNP coordinates with chromatin loops using the BEDTools v2.26.0 pairtobed command [34].

### Dual-luciferase reporter assays for candidate enhancer variants

Transcription factor binding sites affected by candidate SNPs were predicted using JASPAR [37]. Ten SNPs located in enhancer regions and predicted to affect transcription factor binding were selected for dual-luciferase reporter assays. Two genomic regions without predicted regulatory activity were included as negative controls. Approximately 800 bp genomic fragments centered on each candidate SNP and carrying alternative alleles were synthesized or amplified, yielding 24 sequences in total, including controls. Enhancer fragments were amplified from Yorkshire pig genomic DNA using Phanta Max Super-Fidelity DNA Polymerase (Vazyme, Cat. no. P505) and inserted into the pGL3-promoter vector (Promega, Cat. no. E1761) using NcoI (Thermo Fisher Scientific, Cat. no. FD0574) and KpnI (Thermo Fisher Scientific, Cat. no. FD0524). All constructs were verified by Sanger sequencing. Porcine kidney-15 (PK15) cells were cultured at 37 degrees C in 5% CO2. Cells were seeded into 48-well plates and co-transfected with allele-specific reporter constructs and the pRL-TK Renilla luciferase control vector (Promega, Cat. no. E2241) using Lipofectamine 2000 (Invitrogen, Cat. no. 11668500). After 24 h, cells were lysed and luciferase activity was measured using the Dual-Luciferase Reporter Assay System (Yeasen, Cat. no. 11402ES60). Firefly luciferase activity was normalized to Renilla luciferase activity. Each construct was tested in at least three independent biological replicates.

### Electrophoretic mobility shift assay

Nuclear extracts were prepared from PK15 cells using NE-PER Nuclear and Cytoplasmic Extraction Reagents (Thermo Fisher Scientific, Cat. no. 78833) and quantified with a BCA assay (Beyotime, Cat. no. P0010S). Biotin-labeled and unlabeled oligonucleotide probes were synthesized for allele-specific EMSA. For rs334769214, the forward TT probe was 5-prime-TTGTTTTGTTTTGTTTTTTTTTTTTGTCTTT-3-prime, and the forward GG probe was 5-prime-TTGTTTTGTTTTGTTGTTTTTTTTTGTCTTT-3-prime. Binding reactions were performed using the LightShift Chemiluminescent EMSA Kit (Thermo Fisher Scientific, Cat. no. 89880) according to the manufacturer’s protocol. Nuclear protein, biotin-labeled probes and unlabeled competitor probes were combined as indicated for each lane. Reactions were separated on a 6% polyacrylamide gel in 0.5x TBE buffer and transferred at 380 mA for 60 min. Biotin-labeled DNA-protein complexes were detected by chemiluminescence.

### CRISPR activation of the candidate enhancer

To test whether the identified SNP rs334769214-containing enhancer regulates PMAIP1 expression, CRISPR activation was performed using a dCas9-based activation system [38]. sgRNAs targeting the PMAIP1 enhancer were designed using CRISPR-offinder (http://biootools.com/). The selected sgRNA sequence was cloned into the linearized Lenti-sgR-MS2-VP64 backbone and packaged into lentiviral particles. IPECJ2 cells stably expressing dCas9-p300 were transduced with sgRNA-encoding lentivirus. Three days after infection, cells were selected with 10 micrograms/mL blasticidin and harvested for PMAIP1 expression analysis.

### Transcription factor overexpression and allele-specific reporter assays

To evaluate the regulatory effects of candidate transcription factors on enhancer activity at SNP rs334769214, approximately 400 bp enhancer fragments containing either the T or G allele were cloned upstream of the luciferase gene in pGL3-promoter using NcoI and KpnI. Coding sequences for FOXL1, BRCA1 and FOXG1 were synthesized and inserted into pcDNA3.1(+) expression vectors. All constructs were verified by Sanger sequencing. PK15 cells were maintained in DMEM supplemented with 10% fetal bovine serum at 37 degrees C in 5% CO2. For each transfection, cells in 48-well plates received 200 ng of allele-specific luciferase reporter plasmid, 100 ng of transcription factor expression plasmid or empty vector control, and 20 ng of pRL-TK Renilla luciferase plasmid. Luciferase activity was measured 24h after transfection. Each group included at least three biological replicates and three technical replicates.

### Apoptosis assays

To assess the pro-apoptotic function of PMAIP1, the PMAIP1 coding region was cloned into pcDNA3.1(+), with empty vector as the negative control. IPEC-J2 cells, a non-transformed porcine intestinal epithelial cell line originally derived from the jejunum of a neonatal unsuckled piglet, were used as an in vitro model of small-intestinal epithelium [39]. IPEC-J2 cells were cultured in DMEM/F12 medium containing 10% fetal bovine serum at 37 degrees C in 5% CO2 and transfected at 70-80% confluence using Lipofectamine 3000. Cells were harvested 48 h after transfection. Apoptosis was measured using an Annexin V-IF647/PI Cell Apoptosis Detection Kit (Cat. no. G1514-50T). Cells were washed with phosphate-buffered saline, resuspended in binding buffer, stained with Annexin V-FITC and propidium iodide for 15 min at room temperature in the dark, and analyzed within 1h on a BD Accuri C6 flow cytometer. At least 10,000 events were collected per sample. Flow-cytometry populations were classified as viable (Annexin V-negative/PI-negative), early apoptotic (Annexin V-positive/PI-negative), and late apoptotic/dead (Annexin V-positive/PI-positive). The total Annexin V-positive fraction was calculated as the sum of the early apoptotic and late apoptotic/dead populations. For imaging, cells were fixed in 4% paraformaldehyde 48 h after transfection, permeabilized and co-stained with Annexin V antibody and DAPI according to the kit protocol. Images were acquired by super-resolution laser confocal microscopy.

### Statistical analysis

Statistical analyses for reporter assays, CRISPR activation experiments and apoptosis assays were performed using two-sided Student’s t-test for two-group comparisons or one-way ANOVA for comparisons among three or more groups. P < 0.05 was considered statistically significant unless otherwise stated. For genotype-phenotype validation of rs334769214, 872 individuals were included for ADG and 867 for BF. Least-squares means among GG, TG and TT genotypes were compared using Tukey’s test in R v4.3.1 with the agricolae and lsmeans packages. Pairwise comparisons included GG versus TT, TT versus TG and GG versus TG.

## Results

### Descriptive statistical analysis and genetic parameters

For genomic analyses, a subset of Yorkshire pigs with both phenotypic records and genotype data was used, including 7,972 individuals for the birth-to-100-kg average daily gain (ADG) and 7,959 individuals for backfat thickness (BF) (Fig. 1A). Genotyping was performed using a porcine SNP array, which generated 53,261 genome-wide SNP markers. After quality control, 41,843 and 41,845 high-quality SNPs were retained for ADG and BF, respectively (Fig. 1B). Phenotypic records were available for a larger Yorkshire pig population, comprising 58,759 ADG measurements and 58,736 BF measurements. Detailed summary statistics are presented in Table 1. The corresponding pedigree database contained approximately 640,000 records. Based on this dataset, the heritability estimate for ADG was 0.17 (SE = 0.02), and the heritability estimate for BF was 0.31 (SE = 0.04) (Fig. 1C and 1D). The genetic correlation between ADG and BF was 0.55, indicating a moderate positive genetic relationship between growth rate and backfat deposition.

**Fig. 1.**
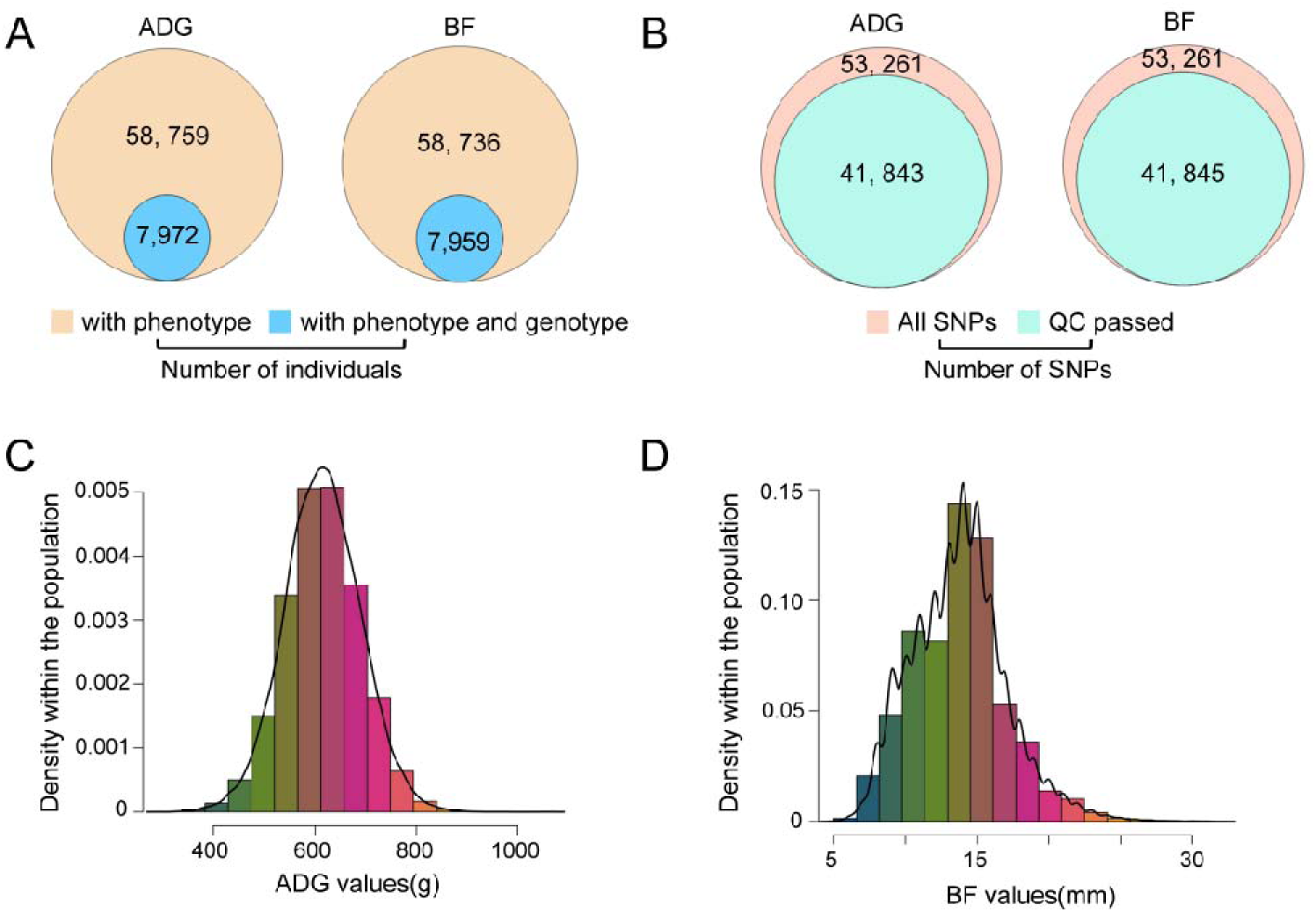
Phenotypic characterization, genetic parameter estimation, and SNP summary for the birth-to-100-kg average daily gain (ADG) and backfat thickness (BF) traits. (A) Summary statistics of phenotype and genotype counts for ADG and BF. (B) Number of SNPs before and after quality control (QC) for ADG and BF. (C) Density distribution of ADG phenotypes, with mean, standard deviation, heritability, and genetic correlation indicated. (D) Density distribution of BF phenotypes are indicated.

**Table 1.** Descriptive statistics for growth and fatness traits in genotyped Yorkshire pigs.

| **Traits** | **n** | **Mean±SD** | **Range** | **HE** |
| --- | --- | --- | --- | --- |
| The birth-to-100-kg<br>ADG (g/day) | 7972 | 623.25±53.89 | 346.70-893.58 | 0.17±0.02 |
| BF (mm) | 7959 | 13.75±3.26 | 5.00-32.00 | 0.31±0.04 |

Note: ADG, average daily gain from birth to 100 kg body weight; BF, backfat thickness; SD, standard deviation; n, number of individuals; HE, heritability estimate; BF was measured between the 10th and 11th ribs at a body weight of $100 \pm 15$ kg.

### GWAS identifies a shared SSC1 locus associated with ADG and BF

To identify genetic variants associated with ADG and BF, we performed GWAS using a linear mixed model. In total, 19 SNPs exceeded the genome-wide significance threshold for ADG and 37 SNPs reached significance for BF. The QQ plots for ADG and BF showed that most observed P-values followed the expected null distribution, indicating adequate control of population stratification and cryptic relatedness, whereas the upward deviation of tail points suggested the presence of trait-associated loci. Detailed annotations, including effect estimates and nearest genes, are provided in Table S1. Among the associated SNPs, 15 ADG-associated and 34 BF-associated variants co-localized within an overlapping interval on SSC1. The ADG-associated SNPs spanned chr1:159,835,711-162,406,821 bp, and the BF-associated SNPs spanned chr1:159,486,739-165,856,655 bp, converging on a shared 2.57 Mb region. Additional trait-specific signals were observed on SSC5, SSC7 and SSC12 for ADG and on SSC18 for BF (Fig. 2). In this study, we focused exclusively on the overlapping genomic region identified on SSC1, as our objective was to elucidate the genetic mechanism associated with both ADG and BF.

**Fig. 2.**
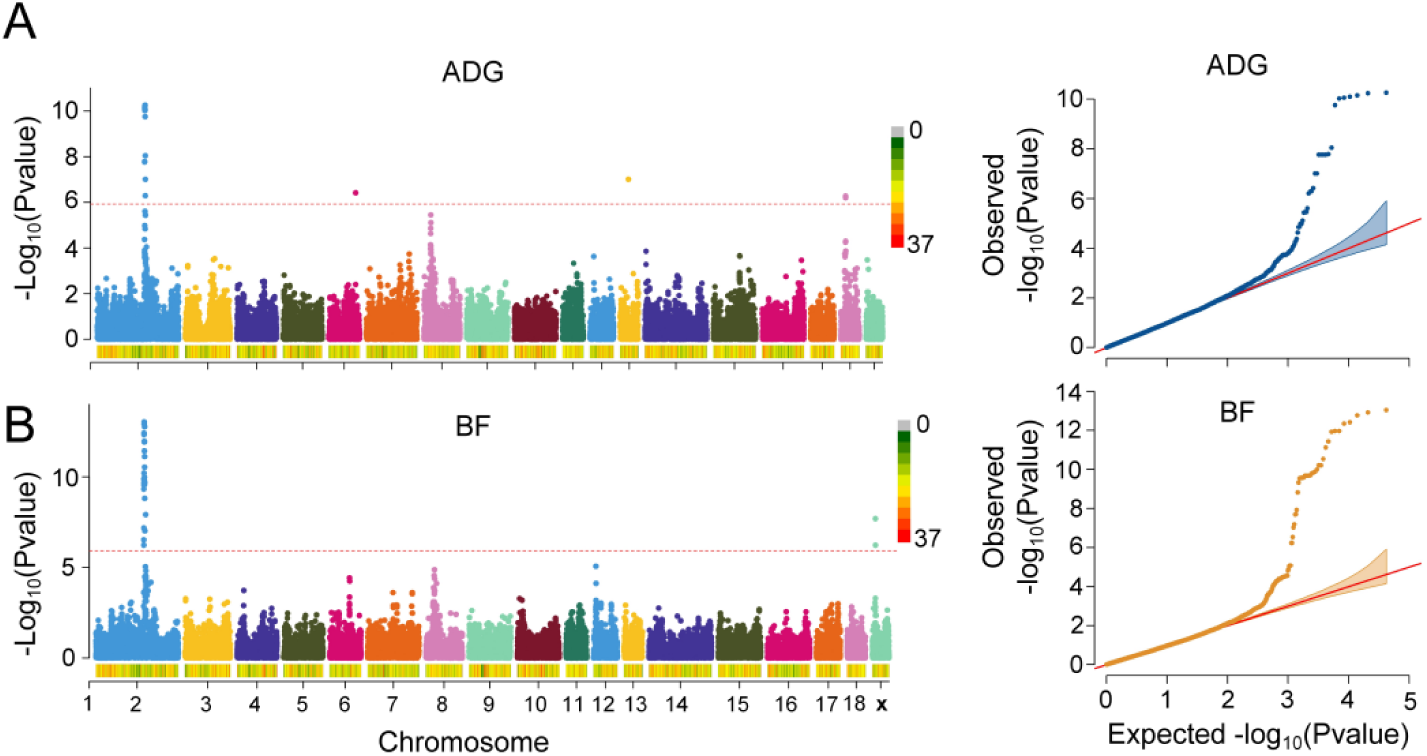
Genome-wide association study of ADG and BF traits in Yorkshire pigs. (A) Manhattan plot for ADG. (B) Manhattan plot for BF. In the Manhattan plots, the red dashed lines indicate the genome-wide significance threshold (−log10(0.05/41845) = 5.92). The QQ plots demonstrate that the observed P-values closely follow the expected null distribution for the majority of SNPs, indicating adequate control of population stratification, while the sharp upward deviation at the right tail indicates association signals for both traits.

### Fine-mapping prioritizes a 1.59-Mb interval within the shared association region

To achieve higher mapping resolution, we examined local Manhattan plots and LD patterns surrounding significant GWAS SNPs on SSC1. By defining the physical boundaries of these signals, we identified a 6.37 Mb candidate region associated with BF (chr1:159,486,739-165,856,655 bp), which encompassed a nested 2.57 Mb interval associated with ADG (chr1:159,835,711-162,406,821 bp) (Fig. 3A and 3B). We subsequently performed LDLA across SSC1 for both traits [21–22]. Using a 95% confidence interval method, LDLA for ADG identified 42 SNPs with the highest likelihood ratio test values spanning the 2.57 Mb region (Fig. 3C), whereas LDLA for BF identified 62 SNPs spanning the broader 6.37 Mb region (Fig. 3D). The peak LRT signals for both traits converged on a tighter shared interval (chr1:160.21-161.80 Mb; 1.59 Mb). Haplotype block analysis of the GWAS-significant region using the confidence interval algorithm in Haploview revealed 10 LD blocks for BF and 3 blocks for ADG. BF-block 3 and ADG-block 2 co-localized within the same 1.59 Mb interval (chr1:160,210,902-161,804,920 bp), prioritizing this region for functional annotation (Fig. S1A). Based on this overlapping interval, we designed targeted capture sequencing for a randomly selected subset of 872 individuals from the original population. Within the targeted 6.37 Mb region sequenced in the 872 individuals, a total of 19,893 SNPs were captured. Among these, 6,891 SNPs were distributed within the 2.57 Mb nested ADG/BF overlapping interval. Association analysis using the high-depth sequencing data revealed that 134 SNPs exceeded the significance threshold for ADG (−log□□ (p) = 5.14), and 167 SNPs reached significance for BF (−log□□(p) = 5.60) (Fig. S1B and S1C). Compared to the original GWAS, targeted sequencing recapitulated the major SSC1 association signal while providing denser variant coverage. Finally, we functionally annotated the 1.59 Mb shared LD block using porcine multi-omics data from small intestine, including H3K27ac, H3K4me3, ATAC-seq and Hi-C (Fig. 3E). A large proportion of trait-associated variants within the interval overlapped transcriptionally active A compartments, and the shared interval was embedded within a topologically associating domain. These data prioritized the 1.59 Mb interval as a regulatory candidate region associated with both ADG and BF.

**Fig 3.**
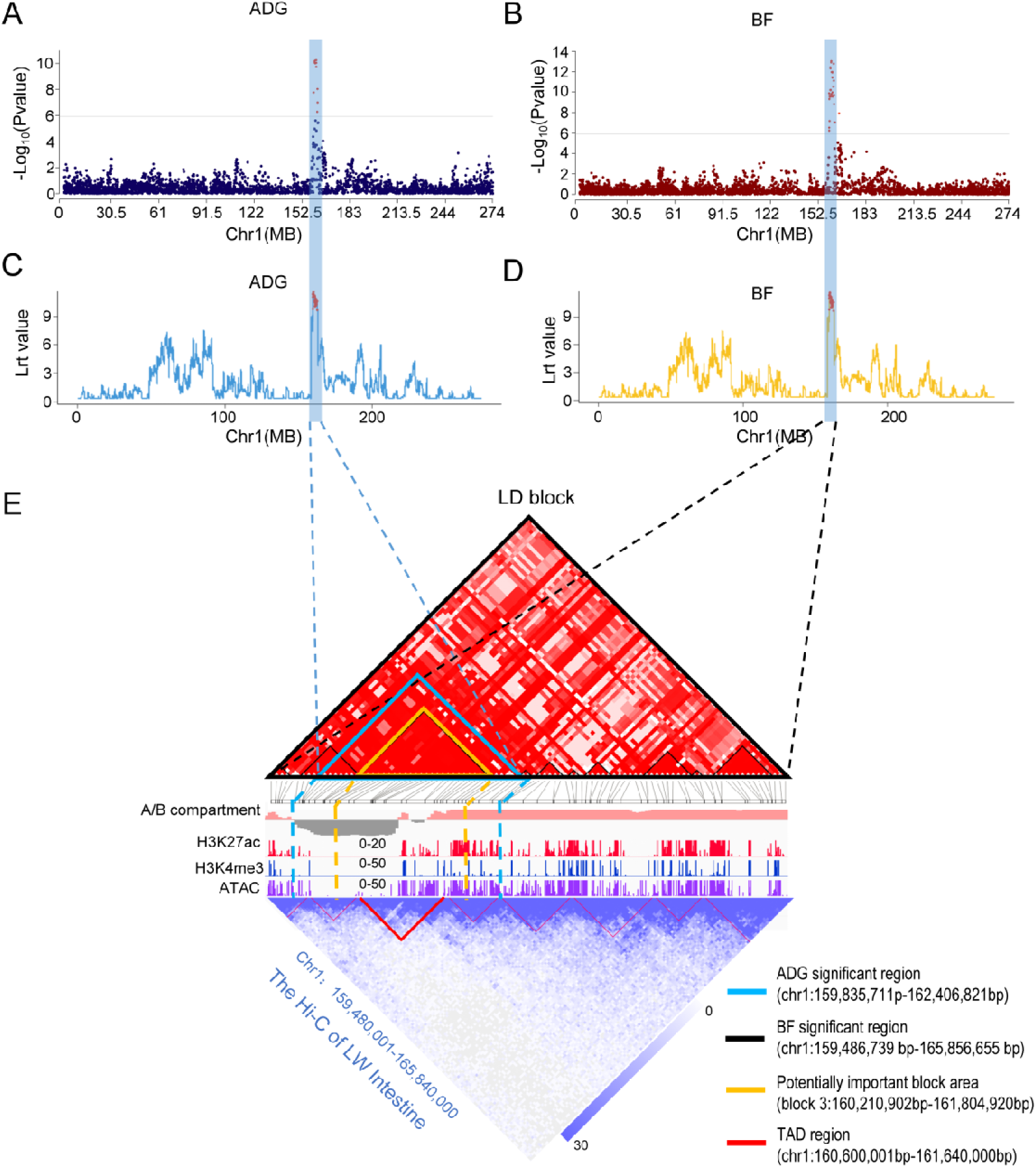
Fine-mapping and multi-omics functional annotation of the shared QTL region on SSC1 associated with ADG and BF. (A, B) Local Manhattan plots showing the association signals on chromosome 1 for ADG (A) and BF (B). The horizontal lines represent the genome-wide significance thresholds, and the two red vertical lines delineate the broad candidate region. (C, D) Linkage disequilibrium locus analysis (LDLA) mapping curves for ADG (C) and BF (D). The y-axis represents the likelihood ratio test (LRT) values, with the peak SNPs highlighted in red. The dashed lines project the peak association signals down to the shared LD block. (E) Integrated epigenomic landscape of the fine-mapped 1.59 Mb shared LD block region. The critical shared interval is highlighted by the yellow boundaries. The tracks display the A/B compartments (active A compartments in pink, inactive B compartments in grey) and epigenetic enrichment profiles (H3K27ac in red, H3K4me3 in light blue, and ATAC-seq signals in purple), which collectively denote active regulatory elements. Hi-C chromatin contact heatmap showed the TAD structure in the fine-mapped 1.59 Mb shared LD block region.

### SNP rs334769214 shows allele-specific enhancer activity and protein binding

As indicated by our haplotype block analysis (Fig. S1A), the significant SNPs identified via targeted capture sequencing are deeply embedded within a shared and tightly linked LD block. Because these variants co-segregate and show highly comparable association signals, association evidence alone could not distinguish the functional variant from tightly linked marker variant. To overcome this limitation and systematically prioritize the regulatory variants associated with both ADG and BF, we applied the IFmut database scoring system [36] to the 134 ADG-associated and 167 BF-associated SNPs identified by capture sequencing. IFmut estimates the regulatory potential of SNPs by integrating their overlap with functional genomic features, including open chromatin regions (OCRs), nucleosome-free regions (NFRs), footprints and transcription factor (TF)-binding motifs (Fig. 4A). Variants classified as levels 1a-3b overlap TF motif sites and at least one class of functional regulatory element, indicating relatively high regulatory potential. Using this criterion, we retained 17 high-confidence candidate SNPs for further analysis, including 6 associated with ADG and 11 associated with BF (Fig. 4B and Table S4).

**Fig. 4.**
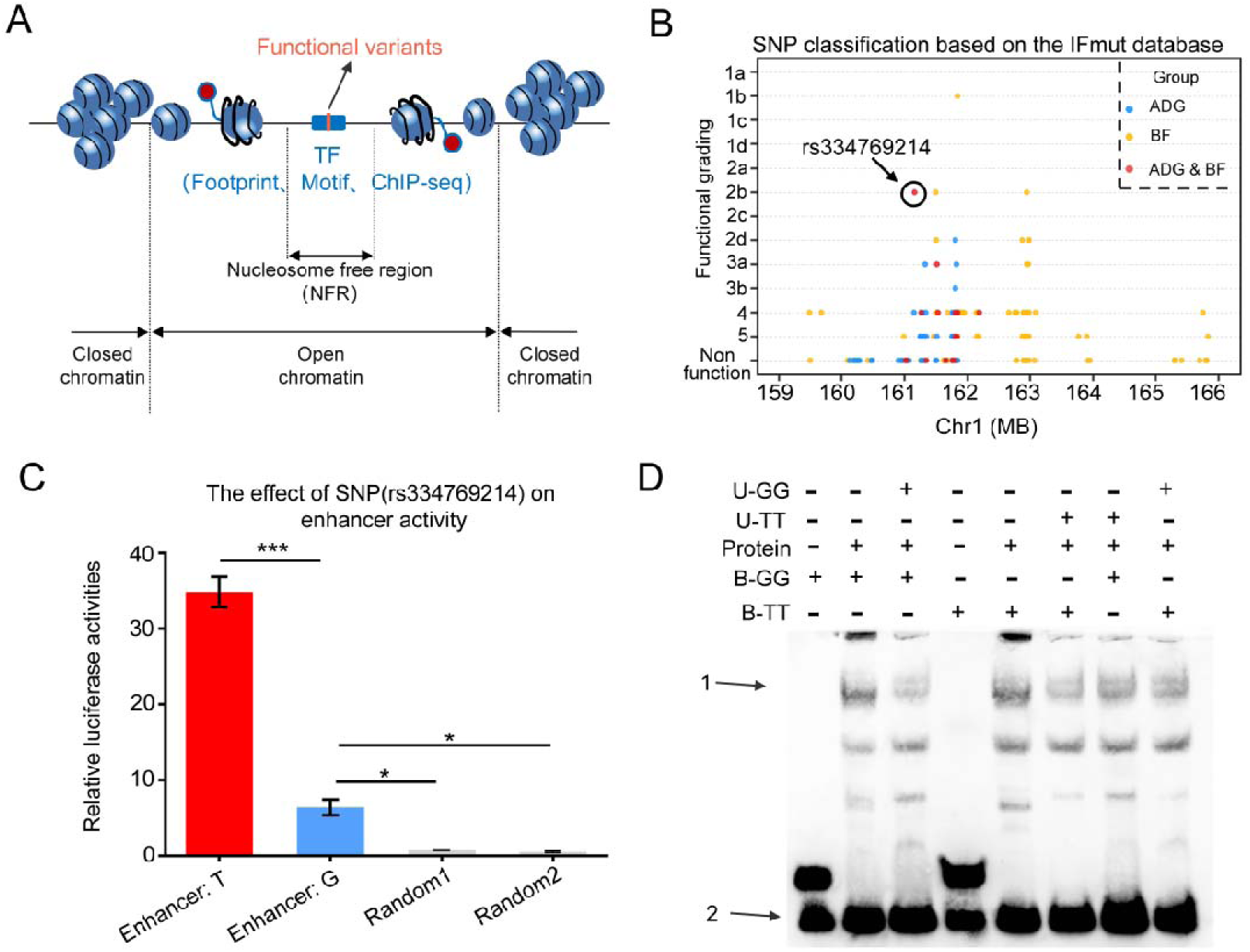
Functional characterization and experimental validation of candidate variants associated with ADG and BF on SSC1. (A) Schematic representation of chromatin accessibility and functional genomic features used to predict transcription factor binding profiles. (B) Genomic distribution and functional priority grading of ADG- and BF-associated variants on chromosome 1 using the IFmut scoring. (C) Dual-luciferase reporter assay validating the allele-specific enhancer activity at the rs334769214 (T/G) locus. R1 and R2 represent two randomly selected, non-regulatory genomic regions used as negative controls. *, P < 0.05; ***, P < 0.001. (D) EMSA validation of allele-specific protein binding at the rs334769214 locus. Arrow 1 indicates the specific protein-DNA complex; Arrow 2 indicates the free probe. “U-GG” and “U-TT” represent unlabeled competitor probes for the GG and TT alleles, respectively. “B-GG” and “B-TT” represent biotin-labeled probes for the GG and TT alleles. “Protein” denotes the addition of nuclear extracts derived from PK15 cells.

Subsequently, we further prioritised candidate SNPs by retaining only those predicted by motifbreakR to have a strong effect on TF-binding motifs and whose corresponding TFs were expressed in the tissues where the associated regulatory elements were active. This filtering strategy yielded 10 significant candidate SNPs, including 4 ADG-associated and 8 BF-associated variants, of which two (rs334769214 and rs332218029) were pleiotropic and shared by both traits (Table 2).

**Table 2.** Using porcine multi-omics data 10 mutation sites identified on the core bases of TF motif*.

| *SNP name* | *Position* | *Alleles* | *location* | *Class* | *TF* | *binding capacity*<sup>1</sup> | *Tissue*<sup>2</sup> |
| --- | --- | --- | --- | --- | --- | --- | --- |
| rs336847218 | 161494072 | T/A | Enhancer | BF 2b | NFIL3 | strong | Liver,Spleen,Pancreas,Lung,Heart,Adipose |
| rs340128203 | 162946582 | G/T | Enhancer | BF 2b | TP53 | strong | Heart,Intestine,Lung,Pancreas |
| rs328524644 | 161842565 | G/T | Enhancer | BF 1b | PRDM4 | strong | Adipose,Muscle |
| rs334769214 | 161160798 | T/G | Enhancer | ADG/BF 2b | BRCA1<br>FOXL1<br>FOXG1 | strong | Cerebellum,Intestine |
| rs336246869 | 161804333 | A/G | Enhancer | ADG 2d | CEBPB | strong | Adipose,Cerebellum,Cerebrum,Cortex,Heart,Intestine,Kidney,Liver,Lung,Muscle,Pancreas,Spleen |
| rs332938092 | 162981595 | G/C | Enhancer | BF 2d | RREB1 | strong | Kidney |
| rs81337512 | 162981627 | A/T | Enhancer | BF 2d | NME1 | strong | Cerebellum |
| rs332218029 | 161516637 | G/A | Enhancer | ADG/BF 3a | MYBL1 | strong | Cerebellum,Muscle |
| rs328468432 | 161329319 | A/G | Enhancer | ADG 3a | ZNF76 | strong | Kidney,Lung,Pancreas,Spleen |
| rs328283673 | 162968089 | C/T | Enhancer | BF 3a | CEBPA | strong | Kidney,Liver |

Note: <sup>1</sup> Combining ability is predicted by the strength of transcription factor binding through the motifbreak R software. <sup>2</sup>The corresponding tissue is predicted by the activity of the control element

To test whether these prioritized variants showed allele-specific regulatory activity, we generated dual-luciferase reporter constructs carrying the corresponding reference or alternative alleles. Reporter assays showed allele-specific enhancer activity for several candidates, supporting the utility of the multi-omics prioritization strategy (Fig. 4C and Fig. S2A). Among the two variants shared between ADG and BF, rs334769214 showed a clear allele-dependent difference in enhancer activity, with the T allele driving higher reporter activity than the G allele (Fig. 4C).

We then performed electrophoretic mobility shift assays (EMSA) with nuclear protein extracts from PK15 cells to assess allele-specific protein binding at rs334769214 and rs332218029. The rs334769214 T-allele probe showed markedly stronger protein binding than the G-allele probe, and competition assays supported the specificity of this interaction (Fig. 4D). In contrast, no clear allele-specific binding difference was detected for rs332218029 (Fig. S2B). Together, these results support rs334769214 as the strongest candidate functional enhancer variant among the shared candidates tested, with allele-specific effects on enhancer activity and protein binding.

### Three-dimensional chromatin interaction and CRISPRa support PMAIP1 as a target of the rs334769214-containing enhancer

To investigate the regulatory role of rs334769214, we used higher-resolution ChIA-PET data generated in-house to link the candidate enhancer to potential target genes. Functional annotation indicated that the enhancer harboring rs334769214 is active in both cerebellum and intestine of Yorkshire pigs (Table S2). Chromatin interactions between the rs334769214-containing enhancer and PMAIP1 were detected in both tissues, supporting PMAIP1 as a candidate target gene of this enhancer (Fig. 5A). Notably, both enhancer activity and PMAIP1 expression were prominent in intestinal tissue, consistent with a potential intestinal regulatory mechanism. To test the functional connection between the rs334769214-containing enhancer and PMAIP1 expression, we used a CRISPRa system (Fig. 5B). Eight sgRNAs targeting the enhancer region were designed and cloned into an sgRNA expression vector. Following lentiviral packaging and transduction into stable p300-expressing IPEC-J2 cells, qPCR analysis showed that sgRNA6 significantly increased PMAIP1 expression relative to controls (Fig. 5C). These results support PMAIP1 as a regulatory target of the rs334769214-associated enhancer.

**Fig. 5.**
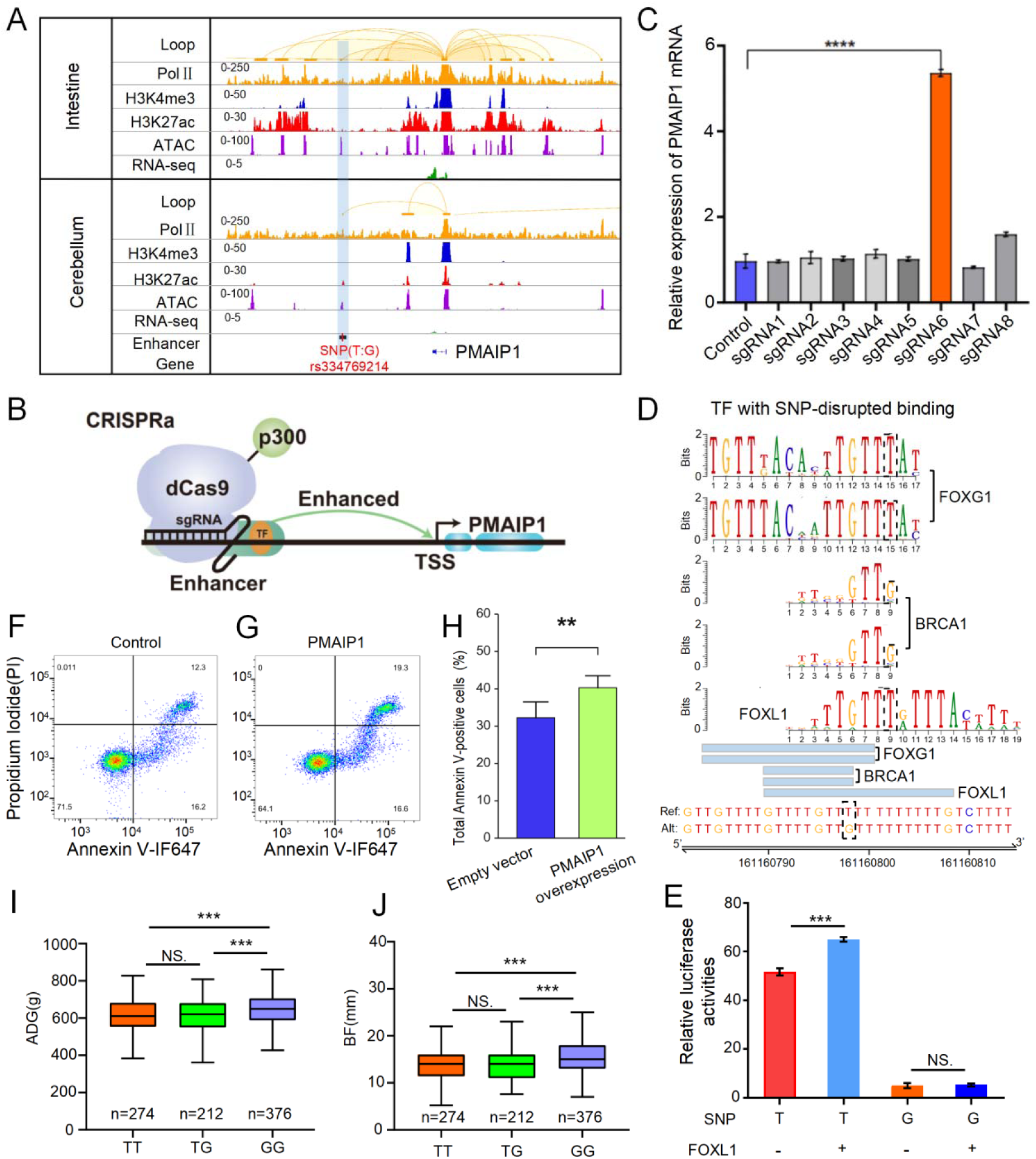
Functional characterization of the rs334769214-associated enhancer and PMAIP1. (A) Chromatin interaction between the rs334769214-associated enhancer region and the target gene PMAIP1. Orange arcs represent chromatin interactions identified by Pol Il ChA-PET, and SNPs regulating PMAIP1 are highlighted in light blue. (B Schematic illustration of the CRISPRa system designed to activate the enhancer region harboring rs334769214. (C) Relative PMAIP1 expression levels detected by qPCR after targeted activation of the enhancer region (sgRNA-targeted) and a negative control region using CRISPRa. Data represent three biological and three technical replicates. ****, P < 0.0001. (D) Prediction of three transcription factors potentially binding to the rs334769214 locus. (E) Dual-luciferase reporter assay evaluating the effects of candidate transcription factors on allele-dependent reporter activity. (F, G) Flow cytometry analysis of apoptosis in IPEC-J2 cells induced by PMAIP1 overexpression at 48 hours post-transfection. (H) Quantification of the total Annexin V-positive fraction across five independent matched experiments. Statistical significance was assessed using a two-sided paired t-test..**, P < 0.01. (I, J) Tukey’s test and multiple comparisons of ADG and BF phenotypic values among different genotypes at the rs334769214 locus across all individuals.

### FOXL1 links rs334769214 to PMAIP1 regulation and intestinal epithelial apoptosis

To further elucidate the regulatory mechanism of rs334769214, we explored transcription factors whose binding may be perturbed by this variant. motifbreakR predicted that the T>G substitution primarily affects motifs for BRCA1, FOXL1 and FOXG1 (Fig. 5D). We then performed dual-luciferase reporter assays in which each transcription factor was co-transfected with reporter constructs carrying either the T or G allele. Among the candidates, only FOXL1 produced an allele-dependent difference in reporter activity: co-transfection of FOXL1 with the T-allele reporter significantly increased enhancer activity compared with the G-allele reporter. These results support a model in which rs334769214 modulates enhancer activity through allele-specific FOXL1 binding (Fig. 5E).

We next assessed the functional consequences of PMAIP1 regulation in IPEC-J2 cells, a non-transformed porcine jejunal epithelial cell model [39]. Across five independent experiments analyzed as matched pairs, the total Annexin V-positive fraction was higher following PMAIP1 overexpression than in empty-vector controls (40.28 ± 7.17% versus 32.25 ± 9.47%; two-sided paired t-test, P = 0.0031; Fig. 5F-5H). In the representative flow cytometry plots, the most apparent redistribution occurred toward the Annexin V-positive/PI-positive population, accompanied by a reduction in the viable Annexin V-negative/PI-negative population. Representative confocal imaging further showed Annexin V-associated staining in PMAIP1-overexpressing cells, providing a qualitative observation consistent with the flow cytometry findings (**Fig. S3**). Together, these observations are consistent with a pro-apoptotic response to PMAIP1 overexpression in IPEC-J2 cells.We then examined the association between rs334769214 genotypes and ADG/BF phenotypes in the capture-sequenced Yorkshire pigs. Individuals carrying the GG genotype showed significantly higher ADG and BF than those carrying the TT genotype (Fig. 5I and 5J). This direction was consistent with the molecular evidence: the T allele exhibited stronger enhancer activity and nuclear-protein binding and was more responsive to FOXL1 overexpression, whereas the G allele represented a relative loss-of-activation state and was associated with higher ADG and BF. Thus, rs334769214 provides a molecular link between allele-specific intestinal enhancer activity, PMAIP1 regulation and coordinated variation in the two traits. Together, these results support rs334769214 as a candidate functional enhancer variant that mediates FOXL1-responsive, allele-dependent regulatory activity and contributes to PMAIP1 regulation, with potential downstream consequences for intestinal epithelial apoptosis and coordinated variation in ADG and BF in pigs.

## Discussion

This study supports a regulatory model linking a shared SSC1 association signal to allele-specific enhancer activity, target-gene regulation and intestinal epithelial apoptosis. By combining GWAS, targeted capture sequencing, LD-based fine-mapping, epigenomic annotation, ChIA-PET and functional assays, we prioritized SNP rs334769214 as a strong candidate functional non-coding variant associated with both ADG and BF. The main functional evidence is consistent with a model in which the T>G substitution influences allele-dependent enhancer activity and the rs334769214-containing region is associated with PMAIP1 regulation.

The direction of the molecular and population-level evidence supports a coherent regulatory model. In our assays, the T allele represented the higher-activity regulatory state, whereas the G allele represented a relative loss-of-activation state at the rs334769214-containing enhancer. This direction is consistent with the known function of PMAIP1/NOXA as a BH3-only pro-apoptotic factor [40]. Because epithelial cell death is closely linked to intestinal epithelial renewal and gut homeostasis [41], and because the intestinal epithelial barrier participates in nutrient handling and systemic metabolic homeostasis [42], attenuation of the FOXL1-PMAIP1-apoptosis axis provides one biologically plausible interpretation of the association between the G allele and higher ADG and BF in this study. This interpretation does not imply that intestinal apoptosis alone determines these traits; rather, it makes explicit the directionally consistent chain from allele-specific enhancer activity, to PMAIP1-associated epithelial apoptosis, to population-level association with growth and fat deposition.

These findings broaden the tissue context for interpreting growth and carcass traits. Previous studies of ADG, BF and related production traits have largely emphasized skeletal muscle and adipose tissue because of their direct roles in growth and fat deposition [13]. The present work highlights the small intestine as an additional regulatory context, consistent with its central role in nutrient digestion, absorption and systemic substrate availability. Thus, the proposed FOXL1-PMAIP1 relationship provides a framework for considering how intestinal epithelial homeostasis may contribute to complex production traits.

Several independent studies further support the biological relevance of the SSC1 region around 160 Mb in pig fatness-related traits. Previous GWAS and meta-GWAS analyses identified signals in this region across diverse genetic backgrounds, including MC4R-centered associations and nearby candidate genes [2,43]. Using sequence-level imputation, Ding et al. narrowed a strong BF association peak to 158.5-162.5 Mb and identified chr1:161511936:T>C as the top imputed SNP, although the regulatory mechanism underlying this signal remained unresolved [44]. Yu et al. subsequently refined a BF QTL to SSC1:160.88-161.51 Mb and nominated PMAIP1-related regulatory mechanisms within this interval [14]. These convergent observations support the relevance of this recurrent SSC1 locus. Building on this genetic evidence, our study connects the region to a coordinated ADG-BF signal and further supports a candidate intestinal regulatory model involving an enhancer variant, allele-specific FOXL1-dependent regulatory activity, three-dimensional enhancer-promoter contact and PMAIP1-associated epithelial apoptosis. Thus, the present work extends previous locus-level associations by providing mechanistic evidence for how regulatory variation in this region may contribute to coordinated variation in growth and fatness.

More broadly, this study demonstrates how post-GWAS interpretation can be strengthened by integrating fine-mapping with regulatory genomics. Targeted capture sequencing identified numerous significant SNPs within a region of strong linkage disequilibrium, underscoring that association statistics alone cannot readily distinguish functional variants from tightly linked markers. In such loci, assigning a GWAS signal to the lead SNP or the nearest annotated gene may overlook distal enhancer-promoter interactions and regulatory variants that are not represented by the strongest association signal [41]. By integrating multi-omics epigenomic annotations, chromatin-interaction maps and experimental validation, we moved beyond the statistical bottleneck and prioritized rs334769214, linking the rs334769214-containing enhancer to PMAIP1 as a candidate target gene and to an allele-specific regulatory mechanism. This framework should be useful for dissecting other complex-trait loci in livestock, while rs334769214 represents a promising functional marker associated with ADG and BF. Its practical value for breeding should be evaluated in the context of the net economic balance between growth rate and carcass fatness.

## Conclusion

In this study, we show the value of integrating GWAS with multi-omics functional characterization for dissecting complex trait loci in livestock. We prioritize rs334769214 as a candidate functional enhancer variant associated with ADG and BF and provide evidence consistent with a model in which the rs334769214-containing enhancer may regulate PMAIP1 through FOXL1-dependent enhancer activity, with potential effects on intestinal epithelial apoptosis. These findings provide mechanistic insight into a non-coding regulatory locus associated with growth and fat deposition traits and nominate rs334769214 as a candidate functional marker for future validation in swine breeding programs.

## Acknowledgements

We sincerely thank Dr. Zhao Shuhong for her invaluable guidance and insightful comments throughout this research. We are grateful to Jing Xu from the Center for Public Instrumentation, College of Animal Science & Technology and College of Veterinary Medicine, Huazhong Agricultural University, for her assistance with confocal microscopy. We also acknowledge the NKLCGI Bioinformatics Platform at Huazhong Agricultural University for providing computational resources.

## Author contributions

H.Y. (Conceptualization [equal], Formal analysis [equal], Investigation [lead], Methodology [lead], Software [equal], Writing - original draft [lead])

M.H. (Formal analysis [lead], Visualization [lead], Software [equal], Writing - original draft [supporting])

Y.X. (Formal analysis [equal], Software [equal], Writing - original draft [supporting])

J.Z. (Investigation [supporting], Methodology [supporting]),

Y.H.Z. (Investigation [supporting], Methodology [supporting])

Y.G. (Software [supporting])

X.D. (Writing - review & editing [supporting])

H.W. (Writing - review & editing [supporting])

X.L. (Funding acquisition [equal],Writing - review & editing [supporting])

Y.X.Z. (Conceptualization [equal], Funding acquisition [equal], Methodology [equal], Project administration [lead], Supervision [lead], Writing - original draft [supporting], Writing - review & editing [supporting])

T.X. (Conceptualization [lead], Funding acquisition [equal], Methodology [equal], Project administration [equal], Supervision [supporting], Writing - original draft [supporting], Writing - review & editing [lead]).

## Supplementary material

Supplementary material accompanies this preprint.

## Data availability

The BioProject record for this study is available at the National Genomics Data Center under accession number PRJCA069169. The processed genotype and phenotype datasets in PLINK format have been deposited in Zenodo under DOI: 10.5281/zenodo.21106985.

---

**Fig. S1.**
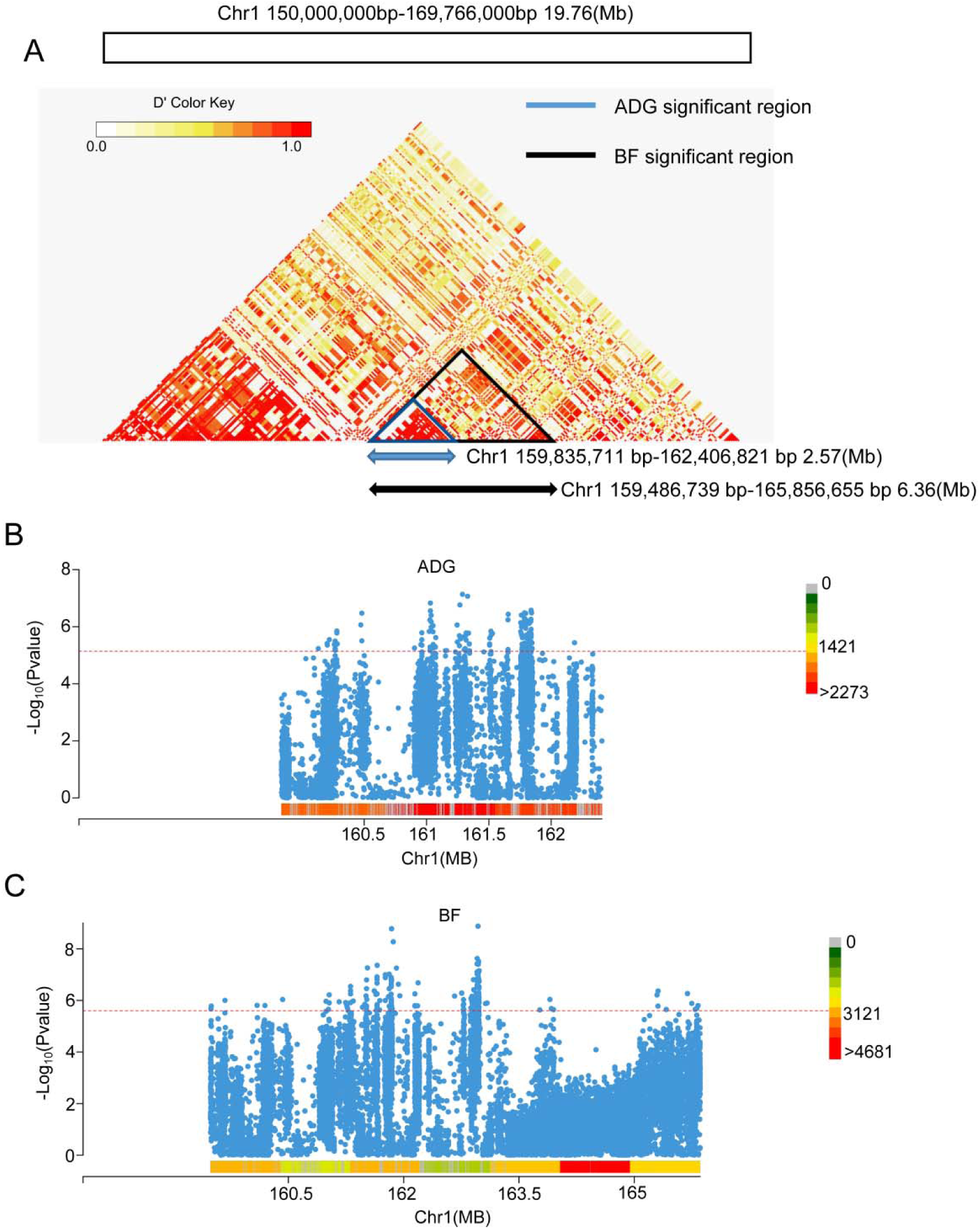
Refinement of the shared SSC1 interval and capture-sequencing association signals for ADG and BF. (A) Haplotype block analysis of the GWAS-significant region on SSC1 using the confidence interval algorithm implemented in Haploview. The peak LRT signals for ADG and BF converged on a shared 1.59 Mb interval (chr1:160,210,902-161,804,920). Within the broader significant regions, 10 LD blocks were identified for BF and 3 for ADG, and BF block 3 overlapped with ADG block 2 within this shared interval, prioritizing the locus for subsequent functional annotation. (B, C) Association analyses based on targeted capture sequencing in 872 randomly selected individuals from the original population. A total of 19,893 SNPs were identified across the targeted 6.37 Mb region, including 6,891 SNPs located within the nested 2.57 Mb overlapping ADG/BF interval. Among these variants, 134 SNPs exceeded the significance threshold for ADG (−log10P = 5.14) (B), and 167 SNPs exceeded the significance threshold for BF (−log10P = 5.60) (C).

**Fig. S2.**
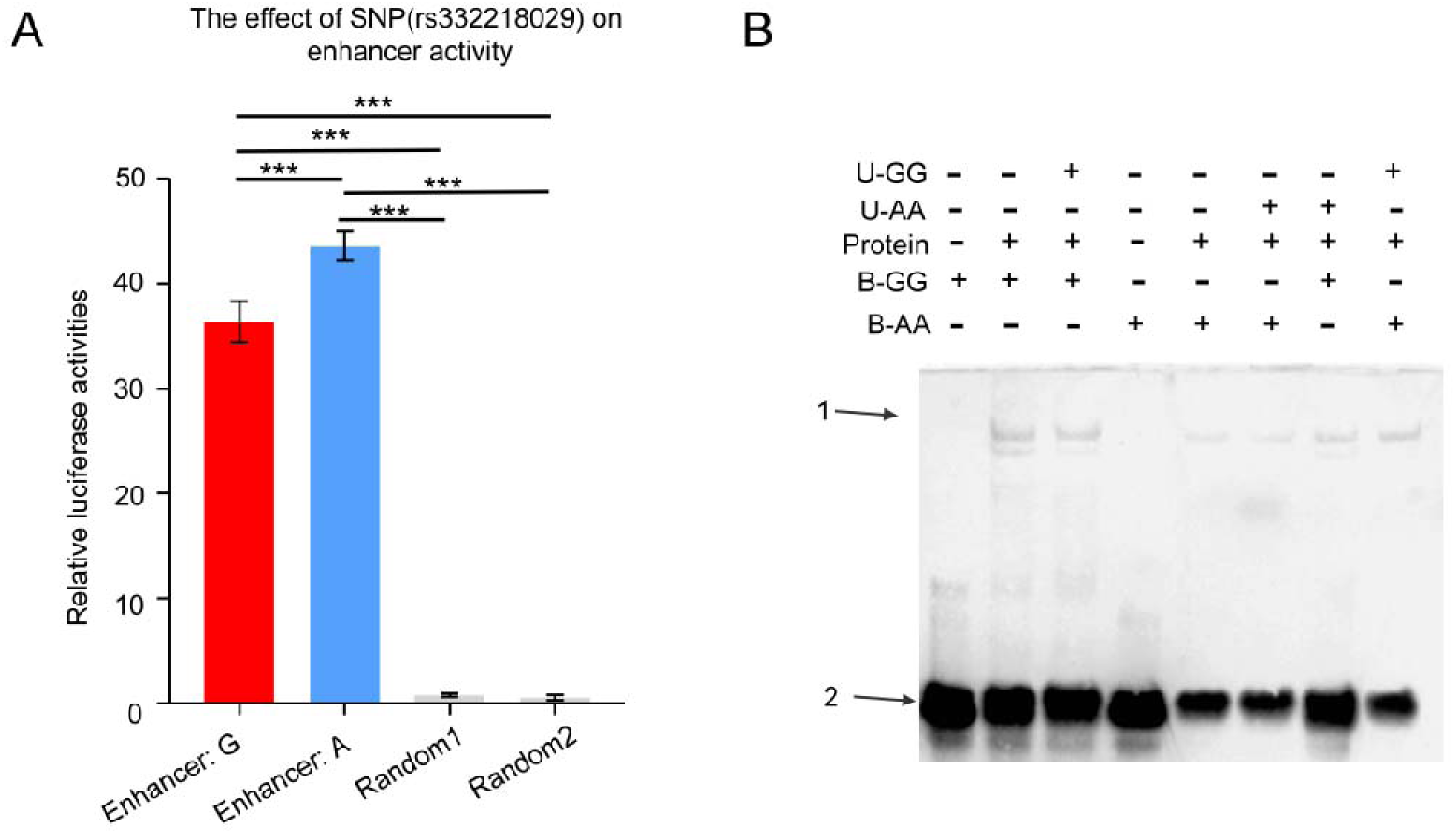
Functional validation of SNP rs332218029. (**A)** Dual-luciferase reporter assay showing the effects of the two alleles of rs332218029 on enhancer activity. Reporter constructs carrying the A allele exhibited significantly higher enhancer activity than those carrying the G allele, whereas the two random control sequences showed minimal activity. Data are presented as mean ± SEM from three independent experiments. Statistical significance was determined by two-sided tests; ***P < 0.001. (**B)** Electrophoretic mobility shift assay (EMSA) for rs332218029. Arrow 1 indicates the specific DNA-protein complex, and arrow 2 indicates the free probe. U-GG and U-AA denote unlabeled competitor probes carrying the G and A alleles, respectively, whereas B-GG and B-AA denote biotin-labeled probes. “Protein” indicates nuclear extracts from PK15 cells.

**Fig. S3.**
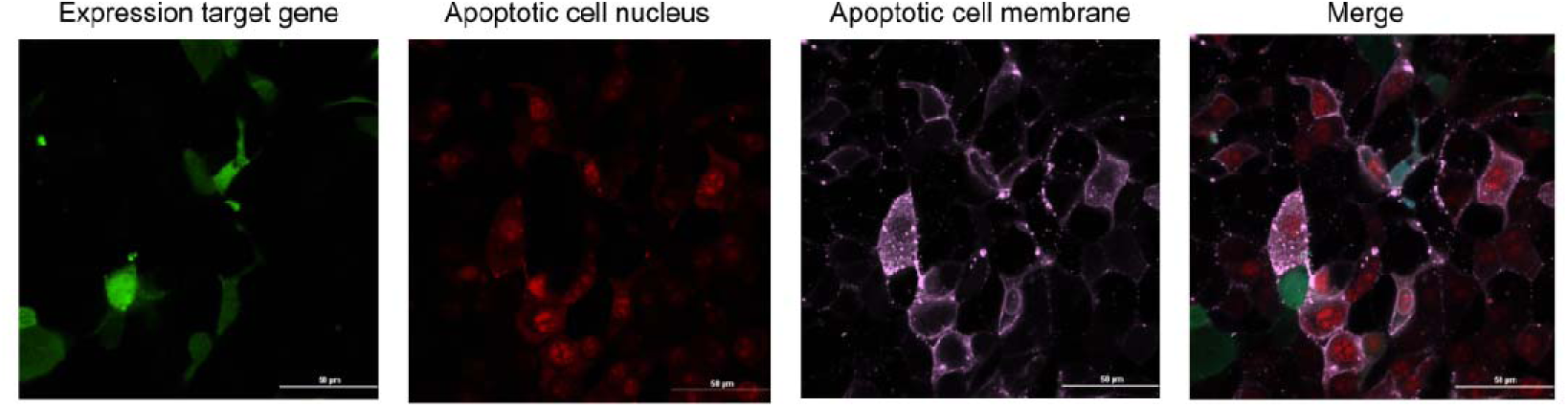
Representative confocal imaging of apoptosis after PMAIP1 overexpression in the porcine intestinal epithelial cell line IPEC-J2. Apoptotic status was assessed 48 h after PMAIP1 overexpression by Annexin V staining. Representative super-resolution confocal images show PMAIP1-expressing cells in green, apoptotic nuclei in red, and apoptotic cell membranes in pink. The merged image illustrates the coexistence of these signals in IPEC-J2 cells and is consistent with different apoptotic states following PMAIP1 overexpression.

**Table S1.** Description of gene and Significant SNPs associated with ADG and BF in Yorkshire Pigs

**Table S2.** Summary of results for linkage disequilibrium and linkage analysis (LDLA).

**Table S3.** Associated SNPs identified by Capture sequencing GWAS analysis on ADG and BF in Yorkshire Pigs

**Table S4.** Screening results of SNP by the functional scoring on ADG and BF in Yorkshire pigs

## References

[1] . Cai Z, Christensen OF, Lund MS, Ostersen T, Sahana G. Large-scale association study on daily weight gain in pigs reveals overlap of genetic factors for growth in humans. BMC Genomics 2022;23:133. 10.1186/s12864-022-08373-3

[2] . Gozalo-Marcilla M, Buntjer J, Johnsson M, et al. Genetic architecture and major genes for backfat thickness in pig lines of diverse genetic backgrounds. Genet Sel Evol 2021;53:76. 10.1186/s12711-021-00671-w

[3] . Jiao S, Maltecca C, Gray KA, Cassady JP. Feed intake, average daily gain, feed efficiency, and real-time ultrasound traits in Duroc pigs: I. Genetic parameter estimation and accuracy of genomic prediction. J Anim Sci 2014;92:2377–2386. 10.2527/jas.2013-7338

[4] . Clutter AC, Brascamp EW. Genetics of performance traits. In: Rothschild MF, Ruvinsky A, editors. The Genetics of the Pig. CAB International; 1998. p. 427–462.

[5] . Ogawa S, Takahashi H, Satoh M. Genetic parameter estimation for pork production and litter performance traits of Landrace, Large White, and Duroc pigs in Japan. J Anim Breed Genet 2023;140:626–638. 10.1111/jbg.12764

[6] . Visscher PM, Wray NR, Zhang Q, et al. 10 years of GWAS discovery: biology, function, and translation. Am J Hum Genet 2017;101:5–22. 10.1016/j.ajhg.2017.06.005

[7] . Hu ZL, Park CA, Reecy JM. (2022). Bringing the Animal QTLdb and CorrDB into the future: meeting new challenges and providing integrated services. Nucleic Acids Res, 50(D1): D956–D961.

[8] . Aherrahrou R, Kaikkonen MU. Technological advancements in functional interpretation of genome-wide association studies (GWAS) findings: bridging the gap to clinical translation. FEBS Lett 2024;598:2852–3. 10.1002/1873-3468.14884

[9] . Kern C, Wang Y, Xu X, Pan Z, Halstead M, Chanthavixay G, et al. Functional annotations of three domestic animal genomes provide vital resources for comparative and agricultural research. Nat Commun 2021;12:1821. 10.1038/s41467-021-22100-8

[10] . Pan Z, Yao Y, Yin H, Cai Z, Wang Y, Bai L, et al. Pig genome functional annotation enhances the biological interpretation of complex traits and human disease. Nat Commun 2021;12:5848. 10.1038/s41467-021-26153-7

[11] . Zhao Y, Hou Y, Xu Y, Luan Y, Zhou H, Qi X, Hu M, Wang D, Wang Z, Fu Y, et al. A compendium and comparative epigenomics analysis of cis-regulatory elements in the pig genome. Nat Commun 2021;12:2217. 10.1038/s41467-021-22448-x

[12] . Teng J, Gao Y, Yin H, Bai Z, Liu S, Zeng H, et al. A compendium of genetic regulatory effects across pig tissues. Nat Genet 2024;56:112–23. 10.1038/s41588-023-01585-7

[13] . Xu Z, Lin Q, Cai X, Zhong Z, Teng J, Li B, et al. Integrating large-scale meta-GWAS and PigGTEx resources to decipher the genetic basis of 232 complex traits in pigs. Natl Sci Rev 2025;12:nwaf048. 10.1093/nsr/nwaf048

[14] . Yu N, Cui D, Li C, Yang S, Qiao C, Xie L. Multi-omics integration reveals Chr1 associated QTL mediating backfat thickness in pigs. J Anim Sci Biotechnol 2025;16:133. 10.1186/s40104-025-01254-1

[15] . Mei Q, Fu C, Sahana G, Chen Y, Yin L, Miao Y, Zhao S, Xiang T. Identification of new semen trait-related candidate genes in Duroc boars through genome-wide association and weighted gene co-expression network analyses. J Anim Sci 2021;99:skab188. 10.1093/jas/skab188

[16] . Jensen J, Christensen OF, Sahana G. DMU - A Package for Analyzing Multivariate Mixed Models in quantitative Genetics and Genomics. In: Proceedings of the 10th World Congress of Genetics Applied to Livestock Production; 2014 Aug 17-22; Vancouver, Canada. https://www.asas.org/docs/default-source/wcgalp-posters/699_paper_9580_manuscript_758_0.pdf?sfvrsn=D;2

[17] . Legarra A, Aguilar I, Misztal I. A relationship matrix including full pedigree and genomic information. J Dairy Sci 2009;92(9):4656–4663. 10.3168/jds.2009-2061

[18] . Aguilar I, Misztal I, Johnson DL, Legarra A, Tsuruta S, Lawlor TJ. Hot topic: A unified approach to utilize phenotypic, full pedigree, and genomic information for genetic evaluation of Holstein final score. J Dairy Sci 2010;93(2):743–752. 10.3168/jds.2009-2730

[19] . Yin L, Zhang H, Tang Z, Xu J, Yin D, Zhang Z, Yuan X, Zhu M, Zhao S, Li X, Liu X. rMVP: A Memory-efficient, Visualization-enhanced, and Parallel-accelerated Tool for Genome-wide Association Study. Genomics Proteomics Bioinformatics 2021;19(4):619–628. 10.1016/j.gpb.2020.10.007

[20] . Barrett JC, Fry B, Maller J, Daly MJ. Haploview: analysis and visualization of LD and haplotype maps. Bioinformatics 2005;21:263–5. 10.1093/bioinformatics/bth457

[21] . Legarra A, Fernando RL. Linear models for joint association and linkage QTL mapping. Genet Sel Evol 2009;41:43. 10.1186/1297-9686-41-43

[22] . Grindflek E, Lien S, Hamland H, Hansen MHS, Kent M, van Son M, et al. Large scale genome-wide association and LDLA mapping study identifies QTLs for boar taint and related sex steroids. BMC Genomics 2011;12:362. 10.1186/1471-2164-12-362

[23] . Druet T, Georges M. A hidden Markov model combining linkage and linkage disequilibrium information for haplotype reconstruction and quantitative trait locus fine mapping. Genetics 2010;184:789–98. 10.1534/genetics.109.108431

[24] . Sartelet A, Druet T, Michaux C, Fasquelle C, Geron S, Tamma N, et al. A splice site variant in the bovine RNF11 gene compromises growth and regulation of the inflammatory response. PLoS Genet 2012;8:e1002581. 10.1371/journal.pgen.1002581

[25] . Garcia-Gamez E, Gutierrez-Gil B, Sanchez JP, Arranz JJ. Replication and refinement of a quantitative trait locus influencing milk protein percentage on ovine chromosome 3. Anim Genet 2012;43:636–41. 10.1111/j.1365-2052.2011.02294.x

[26] . Lander ES, Botstein D. Mapping Mendelian factors underlying quantitative traits using RFLP linkage maps. Genetics 1989;121:185–99. 10.1093/genetics/121.1.185

[27] . Miao, Y., Zhao, Y., Wan, S., Mei, Q., Wang, H., Fu, C., Li, X., Zhao, S., Xu, X. and Xiang, T. (2023) Integrated analysis of genome-wide association studies and 3D epigenomic characteristics reveal the BMP2 gene regulating loin muscle depth in Yorkshire pigs. PLoS Genet., 19, e1010820. 10.1371/journal.pgen.1010820

[28] . Li H, Durbin R. Fast and accurate short read alignment with Burrows-Wheeler transform. Bioinformatics 2009;25(14):1754–1760. 10.1093/bioinformatics/btp324

[29] . McKenna A, Hanna M, Banks E, et al. The Genome Analysis Toolkit: a MapReduce framework for analyzing next-generation DNA sequencing data. Genome Res 2010;20(9):1297–1303. 10.1101/gr.107524.110

[30] . Yang J, Zaitlen NA, Goddard ME, Visscher PM, Price AL. Advantages and pitfalls in the application of mixed-model association methods. Nat Genet 2014;46(2):100–106. 10.1038/ng.2876

[31] . Servant N, Varoquaux N, Lajoie BR, et al. HiC-Pro: an optimized and flexible pipeline for Hi-C data processing. Genome Biol 2015;16:259. 10.1186/s13059-015-0831-x

[32] . Crane E, Bian Q, McCord RP, et al. Condensin-driven remodelling of X chromosome topology during dosage compensation. Nature 2015;523(7559):240–244. 10.1038/nature14450

[33] . Durand NC, Robinson JT, Shamim MS, Machol I, Mesirov JP, Lander ES, et al. Juicebox provides a visualization system for Hi-C contact maps with unlimited zoom. Cell Syst 2016;3:99–101. 10.1016/j.cels.2015.07.012

[34] . Quinlan AR, Hall IM. BEDTools: a flexible suite of utilities for comparing genomic features. Bioinformatics 2010;26:841–2. 10.1093/bioinformatics/btq033

[35] . Robinson JT, Thorvaldsdottir H, Turner D, Mesirov JP. igv.js: an embeddable JavaScript implementation of the Integrative Genomics Viewer (IGV). Bioinformatics 2023;39:btac830. 10.1093/bioinformatics/btac830

[36] . Yin H, Yang L, Zhao Q, et al. Multi-dimensional annotation of porcine variants using genomic and epigenomic features in pigs. BMC Biol 2025;23(1):188. 10.1186/s12915-025-02279-8

[37] . Castro-Mondragon JA, Riudavets-Puig R, Rauluseviciute I, Lemma RB, Turchi L, Blanc-Mathieu R, et al. JASPAR 2022: the 9th release of the open-access database of transcription factor binding profiles. Nucleic Acids Res 2022;50:D165–73. 10.1093/nar/gkab1113

[38] . Li K, Liu Y, Cao H, Zhang Y, Gu Z, Liu X, Yu A, Kaphle P, Dickerson KE, Ni M, Xu J. Interrogation of enhancer function by enhancer-targeting CRISPR epigenetic editing. Nat Commun 2020;11:485. 10.1038/s41467-020-14362-5

[39] . Brosnahan AJ, Brown DR. Porcine IPEC-J2 intestinal epithelial cells in microbiological investigations. Vet Microbiol 2012;156:229–37. 10.1016/j.vetmic.2011.10.017

[40] . Oda E, Ohki R, Murasawa H, Nemoto J, Shibue T, Yamashita T et al. Noxa, a BH3-only member of the Bcl-2 family and candidate mediator of p53-induced apoptosis. Science 2000;288:1053–8. 10.1126/science.288.5468.1053

[41] . Blander JM. On cell death in the intestinal epithelium and its impact on gut homeostasis. Curr Opin Gastroenterol 2018;34:413–9. 10.1097/MOG.0000000000000481

[42] . Chelakkot C, Ghim J, Ryu SH. Mechanisms regulating intestinal barrier integrity and its pathological implications. Exp Mol Med 2018;50:1–9. 10.1038/s12276-018-0126-x

[43] . Zeng H, Zhong Z, Xu Z, Teng J, Wei C, Chen Z, et al. Meta-analysis of genome-wide association studies uncovers shared candidate genes across breeds for pig fatness trait. BMC Genomics 2022;23:786. 10.1186/s12864-022-09036-z

[44] . Ding R, Savegnago R, Liu J, Long N, Tan C, Cai G, et al. The SWine IMputation (SWIM) haplotype reference panel enables nucleotide resolution genetic mapping in pigs. Commun Biol 2023;6:577. 10.1038/s42003-023-04933-9

[45] . Günther C, Neumann H, Neurath MF, Becker C. Apoptosis, necrosis and necroptosis in the gut and intestinal homeostasis. Nat Rev Gastroenterol Hepatol 2013;10:529–36. 10.1038/nrgastro.2013.82

[46] . Aoki R, Shoshkes-Carmel M, Gao N, Shin S, May CL, Golson ML et al. Foxl1-expressing mesenchymal cells constitute the intestinal stem cell niche. Cell Mol Gastroenterol Hepatol 2016;2:175–88. 10.1016/j.jcmgh.2015.12.004

[47] . Fadason T, Schierding W, Lumley T, O’Sullivan JM. Chromatin interactions and expression quantitative trait loci reveal genetic drivers of multimorbidities. Nat Commun 2018;9:5198. 10.1038/s41467-018-07692-y

